# Irradiance modulates thermal niche in a previously undescribed low-light and cold-adapted nano-diatom

**DOI:** 10.1101/2020.07.21.210047

**Authors:** Joshua D. Kling, Kyla J. Kelly, Sophia Pei, Tatiana A. Rynearson, David A. Hutchins

## Abstract

Diatoms have well-recognized roles in fixing and exporting carbon and supplying energy to marine ecosystems, but only recently have we begun to explore the diversity and importance of nano- and pico-diatoms. Here we describe a small (~5 μm) diatom from the genus *Chaetoceros* Isolated from a wintertime temperate estuary (2° C, Narragansett Bay, RI), with a unique obligate specialization for low-light environments (< 120 μmol photons m^-2^ sec^-1^). This diatom exhibits a striking interaction between irradiance and thermal responses whereby as temperatures increase, so does its susceptibility to light stress. Historical 18S rRNA amplicon data from our study site show this isolate was abundant throughout a six-year period, and its presence strongly correlates with winter and early spring months when light and temperature are low. Two ASVs matching this isolate had a circumpolar distribution in Tara Polar Ocean Circle samples, indicating its unusual light and temperature requirements are adaptations to life in a cold, dark environment. We expect this isolate’s low light, psychrophilic niche to shrink as future warming-induced stratification increases both light and temperature levels experienced by high latitude marine phytoplankton.

## Introduction

Photosynthesis by marine phytoplankton is responsible for 50% of the global conversion of inorganic CO_2_ to organic biomass. These photosynthetic protists also perform multiple ecosystem services (1–3), making them essential to study in the context of global climate change (4). Current projections suggest a rise in mean sea surface temperature (SST) of 4 °C by 2100 (5) which could have large impacts on the physiology and composition of these microbial photosynthetic communities, and their ability to sequester carbon (6). However, making predictions about warming effects on phytoplankton can be difficult, because of their still relatively underexplored biological diversity. This is especially true of small (< 5 μm) nano and picoplankton, which prior to the advent of next-generation sequencing have been routinely under-sampled by microscopic techniques (7,8). For example, the smallest known diatom genus (*Minidiscus*) was recently shown to be capable of forming dense blooms, and similar very small diatom groups are now recognized as being globally abundant (8).

Although our knowledge of phytoplankton diversity is expanding, it is an open question how much functional thermal diversity exists within this observed phylogenetic diversity. For instance, phytoplankton communities can typically sustain growth well beyond current mean temperatures. However, excursions above historical thermal maximum thresholds can cause major community restructuring (9), and so affect biogeochemistry (10,11). Lineage-specific predictions of temperature responses have often been based on just a handful of cultured model isolates, such as the diatom *Thalassiosira pseudonana*, (12), the coccolithophore *Emiliania huxleyii* (11), and the diazotrophic cyanobacterium *Trichodesmium erythraeum* (13,14). However, the model organism approach to understanding resilience to rising temperatures in marine phytoplankton undoubtedly under-samples the potential range of thermal responses.

Furthermore, lab-derived growth rates and other proxies for fitness can be uncertain predictors of ecological success. *In situ* variations in temperature, light, and nutrients and interspecific interactions (e.g. competitive or trophic interactions) can make it difficult to apply conclusions from lab-based studies to natural communities.

In this study we expand our knowledge of nanoplankton diversity relative to temperature and light, by characterizing a previously unrecognized nanodiatom from a temperate estuary belonging to the genus *Chaetoceros*. We place its physiology and life history into an environmental context by combining laboratory experiments with a wealth of taxonomic and ecological time series data spanning six years. In addition, we explored the Tara Oceans dataset to explore its global distribution. We report that this *Chaetoceros* isolate exhibits a unique physiological relationship between light and temperature, which skews its abundance strongly towards periods of low light and cold temperatures. This wintertime specialist is thus potentially vulnerable to the warmer conditions expected at mid- and high-latitudes with continuing climate change.

## Methods

### Isolation and Culturing

This diatom was isolated from water collected at the Narragansett Bay Time Series (15) site (latitude 41.47, longitude −71.40) in March, 2018. SST was 2 °C at the time of collection. Surface water was prefiltered through a 100 μm mesh to remove large grazers, and then sorted at the University of Rhode Island Graduate School of Oceanography using a BD Influx flow cytometer (San Jose, CA, USA). Cells approximately 5 μm and smaller with chlorophyll a fluorescence were sorted into 96 well plates containing natural seawater amended with nutrients following the recipe for F media diluted to F/20 (16). Wells showing positive growth over time were transferred to new media while gradually increasing nutrients to F/2 concentrations in Aquil artificial seawater (17). Both initial isolates and stock cultures were maintained at 4 °C and 30 μmol photons * m^-2^ * sec^-1^ of cool-white fluorescent light on a 12:12 L:D cycle, and diluted biweekly with fresh medium. Several dozen morphologically identical strains were collected, all with an apparent sensitivity to light (data not shown). A single isolate strain was selected at random and used for all subsequent experiments.

### Temperature and Light Assays

All culture work was done in climate controlled walk-in incubators under cool white fluorescent lights. Light levels were verified with daily measurements using a LI-250A light meter (LI-COR Biosciences, Lincoln, NE, USA). Cultures were kept in triplicate 15ml polystyrene culture vials, and temperatures for all experiments were set using a series of water baths, each with its own thermostat, heater, and cooling element. Replicates were kept in exponential phase by diluting cultures with sterile medium when biomass reached a predetermined threshold. Cultures were acclimated to each combination of irradiance (15, 30, 50, 60, 70, 100, and 120 μmol photons m^-2^ sec^-1^) and temperature (2, 8, 12, 14, 16, 18, 20, 22, 24, 26 °C) for two weeks.

After acclimation, growth rates were determined daily using *in vivo* fluorescence on a Turner AU-10 fluorometer (Turner Designs Inc., Sunnyvale, CA, USA) for an additional seven to ten days. *In vivo* fluorescence was used as a proxy for photosynthetic biomass, because it allowed efficient daily measurements of a large number of simultaneously maintained cultures (as many as 90 at a time). All well-acclimated replicates were kept in the same nutrient conditions, and growth rates were calculated from *in-vivo* fluorescence of each individual replicate measured relative to itself over time (9,18–21). Although *In-vivo* fluorescence was never used to compare biomass across different light and temperature treatments, in a pilot study we observed that fluorescence and cell counts increased linearly even across different light and temperature treatments (Figure S1).

Specific growth rates were calculated with the GrowthTools R package (DOI:10.5281/zenodo.3634918) using the slope of a regression line fit to the log of these data (20). Growth rates for cultures acclimated to 16 °C were calculated for seven light intensities. In addition, we measured growth rates versus each temperature in cultures acclimated to irradiances of 15, 30, or 50 μmol photons m^-2^ sec^-1^. GrowthTools (DOI:10.5281/zenodo.3634918) was used again to calculate thermal performance curves (TPCs) using the Eppley-Norberg model (22,23). For the thermal curve done under 15 μmol photons m^-2^ sec^-1^, 4 °C was used as the lowest temperature instead of 2 °C.

In addition to acclimated growth experiments, we exposed the cultures to short-term doses of extreme light levels on the order of hundreds of μmol photons m^-2^ sec^-1^, similar to published diatom light-stress experiments (24,25). For these experiments we used ~638 μmol photons m^-2^ sec^-1^, which approximated the highest value measured in a 50-year dataset of surface irradiance from the Narragansett Bay Time Series (https://web.uri.edu/gso/research/plankton/data/). Triplicate cultures acclimated to either 4 °C or 16 °C and 30 μmol photons m^-2^ sec^-1^ were exposed to this extreme light level for one, three, or six hours, and compared to triplicate cultures that were not exposed (negative control), or were continuously exposed (positive control). After the exposure period, cultures were moved back to 30 μmol photons m^-2^ sec^-1^ and fluorescence was recorded twice daily over three days of a 12:12 hour L:D cycle.

### Sequencing

For sequencing, 200 ml of dense culture was filtered onto a 0.22 μm polyethersulfone Whatman Nuclepore filter (GE Healthcare, Chicago, IL, USA), flash frozen with liquid nitrogen and stored at −80 °C. DNA was extracted using a DNEasy Power Water kit (Qiagen, German Town, MD, USA), and prepared for sequencing using the Nextera DNA Flex Library Prep kit (Illumina, San Diego, CA, USA). Sequencing was done at the University of Southern California’s Genome Core on a Illumina Nextseq 550. Raw sequence data was quality checked using Fastqc (https://www.bioinformatics.babraham.ac.uk/projects/fastqc/, v. 0.11.8) and Multiqc (26)(v. 1.6) and low quality bases were removed with Trimmomatic (27)(v. 0.38). To recover 18S rRNA gene sequences, we used Bowtie2 (28)(v. 2.3.5) to map all reads to a dataset of 200 complete or nearly complete (>1000 bp) *Chaetoceros* 18S rRNA gene sequences downloaded from NCBI. Reads that mapped even once were recovered using Seqtk (https://github.com/lh3/seqtk, v. 1.3) and assembled with SPAdes (29)(v. 3.11). To identify our isolate, full length copies of the 18S rRNA gene sequence were downloaded from NCBI for 25 distinctly named species of *Chaetoceros*. The pennate diatom *Pseudo-nitzschia australis* was included as an outgroup. All sequences were aligned using Muscle (30)(v. 3.8.31), the alignment trimmed using trimAL (31)(v. 1.4.15), and FastTree (32)(v. 2.1.10) was used to construct a phylogenetic tree.

Six years of amplicon sequencing data from our study site using diatom-specific primers matching the V4 hypervariable region of the 18S rRNA gene (33) were obtained from (15). Raw sequence data were downloaded from NCBI (PRJNA327394) and quality filtered as for the Illumina sequencing. Quality-controlled reads were merged and denoised into Amplicon Sequence Variants (ASVs) using DADA2 (Callahan et al. 2016, v. 1.14.0). BLAST (McGinnis and Madden 2004, v. 2.9.0) was used to identify ASVs that matched the V4 rRNA gene sequence from the full length sequence assembled from our genomic data.

We also mined years of observational data to put the occurrence of our isolate ASV in Narragansett Bay into a long-term temperature and irradiance context. Sea surface temperature (SST) data matching the amplicon sequencing data were downloaded from the Narragansett Bay Time-series website (https://web.uri.edu/gso/research/plankton/data/). Dates without SST measurements from the time-series dataset were supplemented by SST data from the National Data Buoy Centers station QPTR1 – 8454049 at nearby Quonset Point (https://www.ndbc.noaa.gov/station_page.php?station=qptr1). Irradiance data for Narragansett Bay were downloaded from the National Research Reserve System’s Central Data Management Office website (https://cdmo.baruch.sc.edu/dges/) for station NARPCMET. To avoid over-inflating potential correlations between changing relative abundance within the diatom community (caused by differences in 18S rRNA gene copy number) and environmental factors, we used one percent relative abundance as a threshold and calculated the changing probability of an observation being above this threshold under different conditions. For instance, if this diatom had a relative abundance > 1% in half of a group of samples then its probability of detection was 0.5.

We utilized the Tara Oceans V9 amplicon dataset (36) to understand the distribution of this isolate beyond Narragansett Bay. Sequence data from low- and mid-latitudes previously analyzed and resolved into ASVs using DADA2 were screened for the presence of this isolate diatom using BLAST (35,37). In addition, we downloaded the Tara Polar dataset and analyzed all amplicon sequencing data for these Arctic Ocean surface samples collected on filters with a pore size < 5 μm. Reads were denoised following the methods of (37).

### Statistics and Data Availability

All statistics for analyzing these data and graphic visualizations were done using R (38)(v. 3.6.1) and Rstudio (39)(v. 1.13.83). Differences between light treatments were determined using a one-way ANOVA and the Tukey test, while differences between TPCs were assessed using a repeated measure ANOVA. In both cases, significance was determined at the p < 0.05 level. All environmental data compiled in this study, the output from DADA2, scripts used to download the Tara Polar Ocean Circle samples, and scripts used in analysis have been made publicly available at https://figshare.com/projects/nanodiatom_temp_light/74283. Sequence data can be found on NCBI under the SRA accession PRJNA608686 (raw Illumina reads) and MT742785 (assembled 18S rRNA sequence).

## Results

### 18S rRNA resolved taxonomy

Short read sequencing produced nine million 150bp paired-end Illumina reads. Mapping reads to 200 full length *Chaetoceros spp*. 18S sequences and assembling all mapped reads produced a single contig 1812 bp long. When BLASTed against the nt database, excluding all non-cultured isolates, this assembled 18S rRNA gene sequence was the closest match to *C. cf. wighamii* strain BH65_48, with 99.8% identity across 92% of the sequence (accession KY980353.1). Unfortunately, isolation information was not available for this strain; however, the next closest match was to another *C. cf. wighamii* strain from the Roscoff Culture Collection (RCC3008, KT860959.1) at 100% identity across 90% of the query. This strain was isolated from the coastal Baltic Sea in 2010 at 4 °C, and was maintained at 50 μmol photons m^-2^ sec^-1^, similar to our isolate. Aligning 25 full length 18S sequences for named *Chaetoceros* species (Table S1) allowed us to construct a high quality phylogenetic tree (average maximum likelihood = 0.91) of this genus (Figure 1). *C. cf. wighamii* was the closest branching sequence, followed by temperate isolates *C. throndsenii* from the Gulf of Naples (93.6% ID and 96% coverage), and *C. lorenzianus* (94.1% ID and 90% coverage) and *C. constrictus* (93% ID and 94.3% coverage) from Las Cruces, Chile.

**Figure 1:**
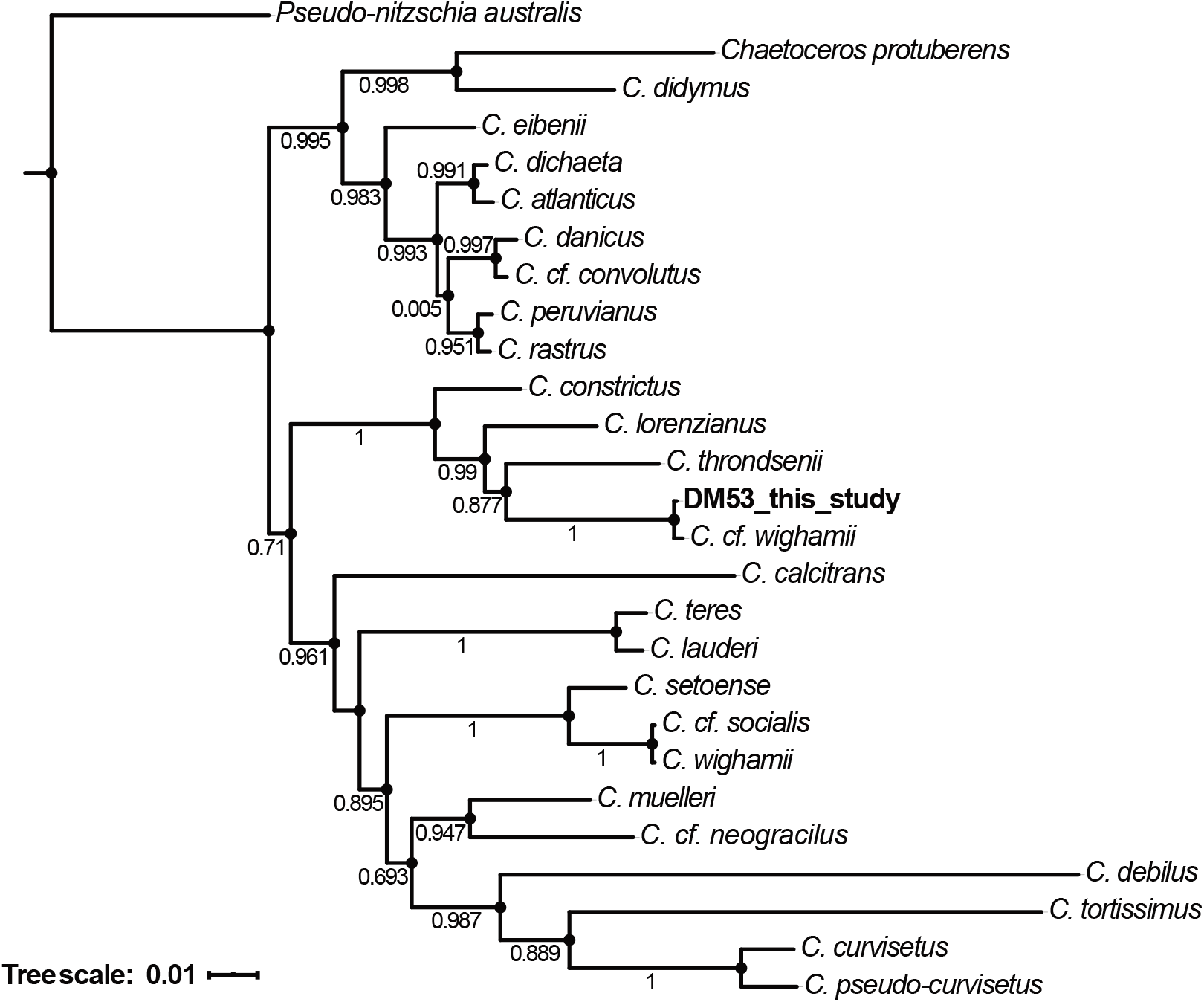
Phylogenetic tree representing the diversity of the diatom genus *Chaetoceros* constructed using full-length 18S rRNA gene sequences from NCBI. A sequence from the pennate diatom species *Pseudo-nitzschia australis* is included as an outgroup. The isolate described in this study, DM53, is highlighted in bold.

### Light Curve

When grown at 16 °C, this diatom isolate had an asymmetric response to increasing light levels, skewed towards low irradiance (Figure 2). Although at the lowest light level tested (15 μmol photons m^-2^ sec^-1^) the specific growth rate was only 0.13 day^-1^ (±0.01), when irradiance was increased to 30 μmol photons m^-2^ sec^-1^, the growth rate nearly tripled, to 0.33 day^-1^ (±0.03). At this light level the specific growth rate was significantly higher than at every other irradiance level tested (*p* < 0.05). Light levels beyond 30 μmol photons m^-2^ sec^-1^ caused the growth rate to rapidly decrease again. For 50, 60, 70 μmol photons m^-2^ sec^-1^ the specific growth rates were 0.19 (±0.04), 0.20 (±0.02), and 0.18 day^-1^ (±0.04) respectively, and they were statistically indistinguishable from each other (*p* > 0.05). Growth rates of our *Chaetoceros* isolate dropped significantly to only 0.10 ±0.01 day^-1^ at 100 μmol photons m^-2^ sec^-1^ (*p* < 0.05), and growth was negative (−0.06 day^-1^, ±0.01) when light levels were increased to 120 μmol photons m^-2^ sec^-1^, leading to eventual cell death (Figure 2).

**Figure 2:**
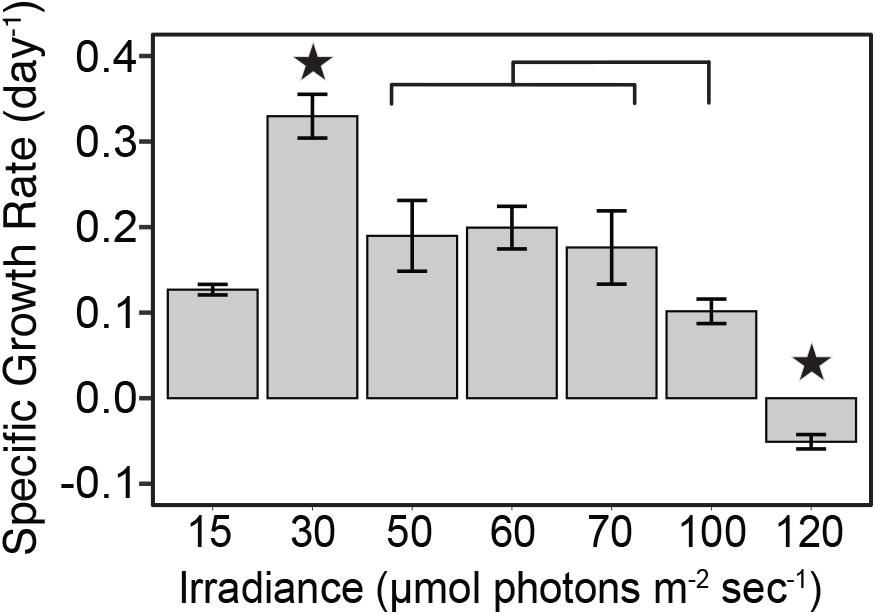
Growth rates at 16 °C across a range of seven light levels for a novel *Chaetoceros sp*. isolate. Error bars represent ±1 standard deviation. Stars show treatments that are statistically significant (*p* < 0.05) compared to all other treatments via one-way ANOVA. Brackets indicate statistical significance between specific samples.

### Thermal Curves

Interactive effects between light and thermal niche for this *Chaetoceros* isolate from the thermal performance curves (TPCs) at three different irradiance levels (15, 30, and 50 μmol photons m^-2^ sec^-1^) are shown in Figure 3a and Table 1. Here we abbreviate the light treatments as low, optimal, and high light. Using a repeat measures ANOVA, each of these TPCs was significantly different from each other (Figure 3a, *p* < 0.001). Comparing values obtained from the Eppley-Norberg model, the full range of growth temperatures (the thermal niche width) was broadest at optimal and low light (27.0 and 25.7 °C, respectively), while the high light niche width was only 23.2 °C (Table 1). The difference in modelled niche width compared to optimal light was manifested as a 1.3 °C decrease in the upper temperature limit (Tmax) under low light (23.7 °C), and as a 3.5 °C increase in the lower temperature limit (Tmin), under high light (1.5 °C) (Table 1). The model predicted that the optimal growth temperature (Topt) would be higher at 30 μmol photons m^-2^ sec^-1^ (17.2 °C ±0.86, Fig 3b, Table 1). The Topt decreased under both low and high light, and although it fell farther in low light, both low and high light Topts were within one standard deviation of each other (13.7 ±0.87 and 15.2 ±1.17 °C respectively). The maximum specific growth rate (μMax) across these TPC models was estimated to be highest under optimal light (0.28 day^-1^ ±0.02) and high light (0.25 day^-1^ ±0.02, Figure 3c, Table 1).

**Figure 3:**
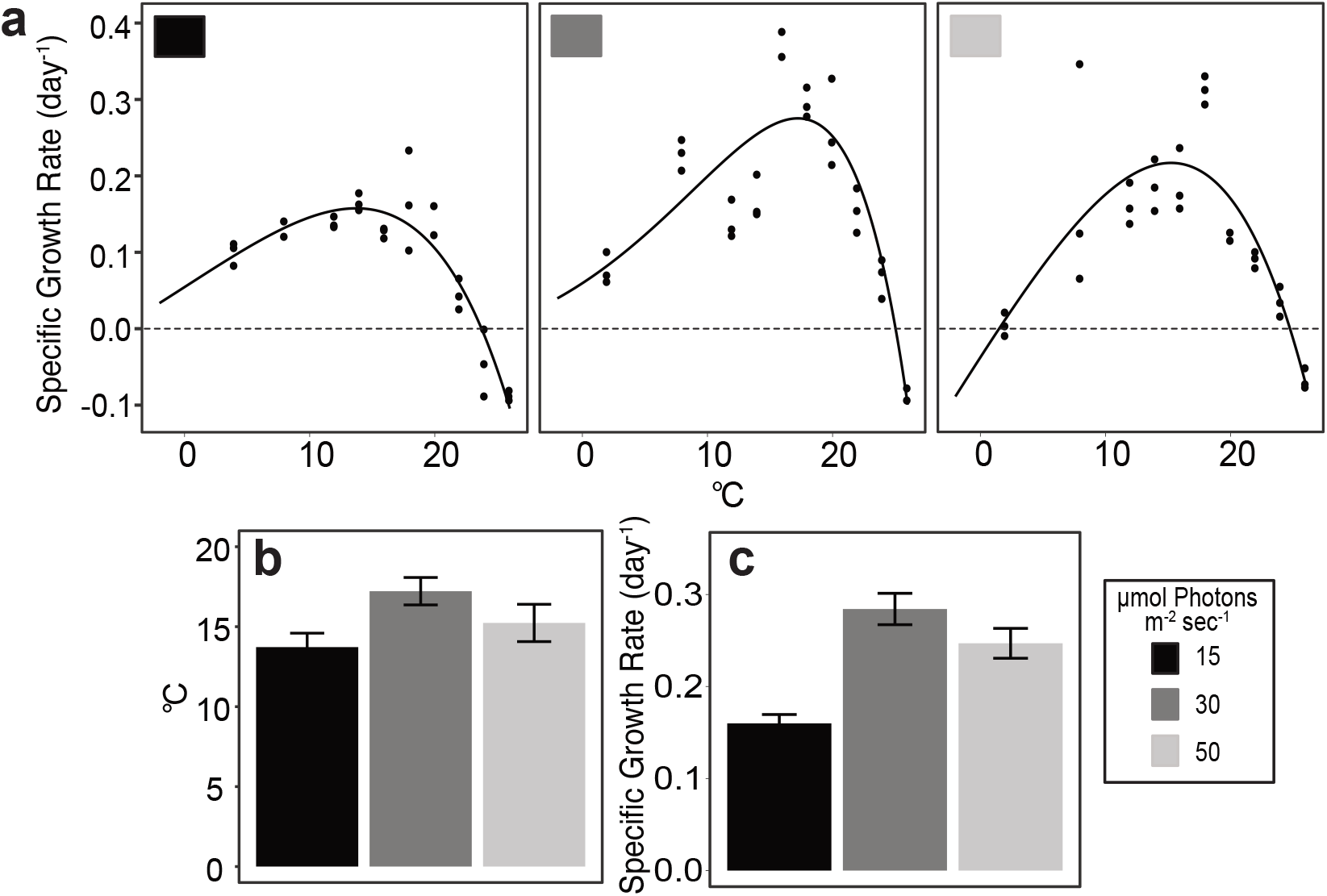
**a)** Thermal performance curves across three light levels. For all best-fit curves, r^2^ > 0.7 and all are significantly different from each other using repeat measure ANOVA (*p* < 0.001). Differences between light treatments are shown for the **b**) thermal optimum (Topt) and **c**) maximum growth rate (μMax). Error bars show ±1 standard deviation within the modelled optimal temperatures and growth rates for each.

**Table 1:**
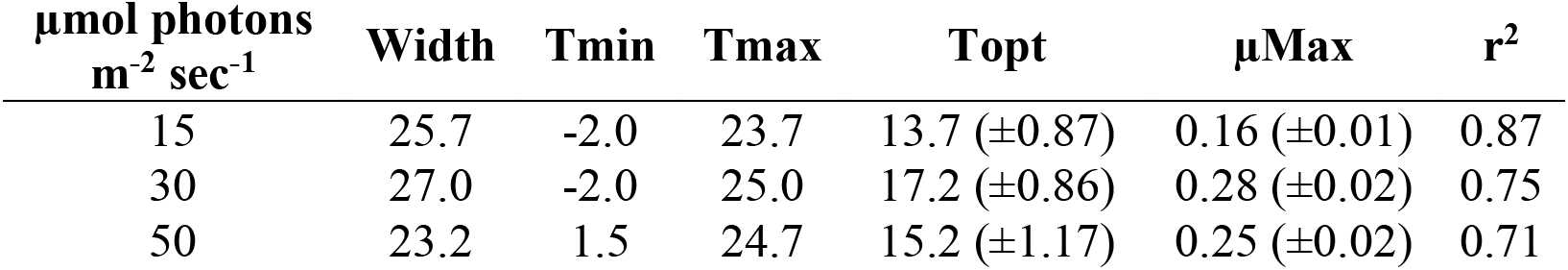
Thermal performance curve (TPC) parameters calculated at three light intensities. Standard deviations are shown where available, within parenthesis.

| $\mu\text{mol photons m}^{-2} \text{ sec}^{-1}$ | Width | Tmin | Tmax | Topt | $\mu\text{Max}$ | $r^2$ |
| --- | --- | --- | --- | --- | --- | --- |
| 15 | 25.7 | -2.0 | 23.7 | 13.7 ($\pm 0.87$) | 0.16 ($\pm 0.01$) | 0.87 |
| 30 | 27.0 | -2.0 | 25.0 | 17.2 ($\pm 0.86$) | 0.28 ($\pm 0.02$) | 0.75 |
| 50 | 23.2 | 1.5 | 24.7 | 15.2 ($\pm 1.17$) | 0.25 ($\pm 0.02$) | 0.71 |

### Response to Extreme Light Stress

In addition to considering how this diatom responded when acclimated to constant light and temperature conditions, we also assessed how it responded to pulses of extreme light at 4 and 16 °C. At both temperatures, constant exposure to extreme light levels (638 μmol photons m^-2^ sec^-1^) was lethal. At 4 °C the constant extreme light treatment (positive control) fluorescence decreased steadily until it reached the lower limit of detection at the very end of this experiment (Figure 4a); however, at 16 °C fluorescence reached the lower limit after just 24 hours (Figure 4c), 3.3x faster (Figure 4b & d). At the lower temperature this *Chaetoceros* isolate maintained positive growth even after being exposed to extreme light for six hours, although exposure for both three and six hours significantly decreased the growth rate compared to cultures never exposed to extreme light (negative control, *p* < 0.05). Similarly, measured growth rates were significantly lower after three- and six-hour exposures compared with the negative control. Cells acclimated to 16 °C also had significantly lower growth rates compared to unexposed cultures (*p* < 0.05); however, unlike at the colder temperature, exposure to extreme light for six hours was lethal.

**Figure 4:**
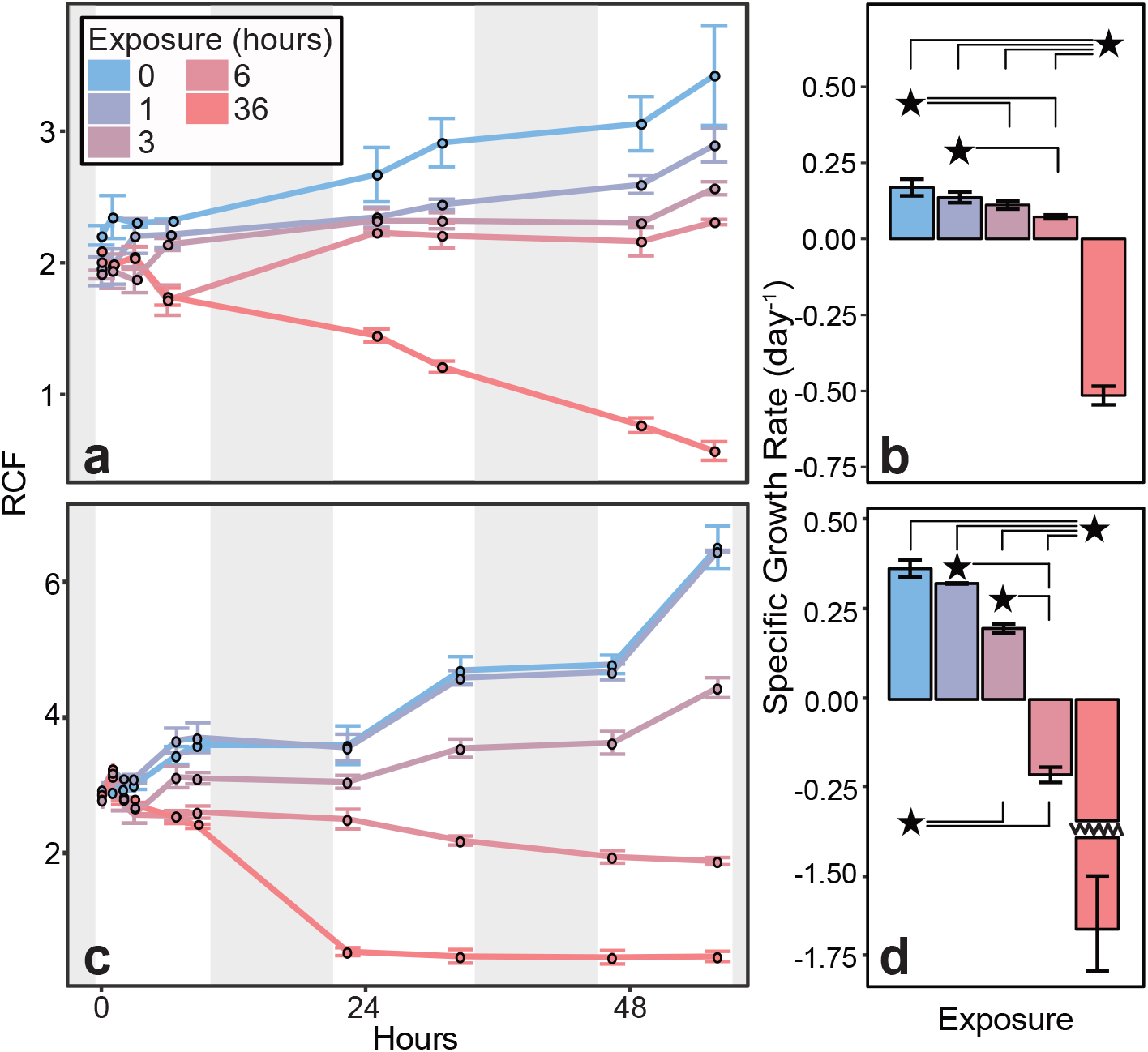
Effects of temperature and light exposure time on cultures of the novel *Chaetoceros sp*. isolate exposed to 638 μmol photons m^-2^ sec^-1^, the highest incident light level recorded in 41 years of data from its isolation location. **a)** Fluorescence (RCF) and **b)** specific growth rates (d^-1^) of cultures grown at 4 °C under extreme light exposure. **c)** and **d)** show the same parameters for cultures grown under extreme light at 16 °C. In panels **a** & **c,** periods of darkness are shown as grey bands. **b** & **d** depict the growth rates for each exposure treatment after three days. All error bars indicate ±1 standard deviation. Stars and brackets show treatments that are significantly different by one-way ANOVA (*p* < 0.05).

### Environmental Amplicon Data

Before using the assembled 18S rRNA sequence to look for this diatom in available amplicon sequencing data from Narragansett Bay, we first confirmed that there was enough diversity within the V4 hypervariable region across this genus to distinguish our isolate from other *Chaetoceros spp.*. Using aligned V4 regions of the same sequences in Figure 1 we were able to construct a phylogenetic tree (Figure S2) that separated our isolate from other members of the genus. Processing sequencing data from Narragansett Bay resulted in 5170 distinct amplicon sequence variants (ASVs); however, only 20 of these had an average relative abundance greater than one percent of recovered amplicons across the data set. When BLASTed, the most common diatom genera were *Thalassiosira* (eight), *Skeletonema* (four), and *Minidiscus* (two)(Table S2). These are consistent with previous observations at Narragansett Bay, where *Thalassiosira* and *Skeletonema* often dominate the diatom community (15,40). Of these 20 most abundant recovered ASVs, one was a perfect match (100% ID and 100% coverage) to the 18S rRNA gene sequence of our isolate.

This ASV was greater than one percent of the total recovered amplicons in 28 of the 80 samples. It had a ~0.5 probability of detection in samples in January, February, and March (Figure 5a). This probability peaked at 0.83 in April, before decreasing to 0.43 again in May. In June through August the probability of detection was ~0.2 and dropped to zero in September and October, before rising again to 0.5 in November and December (Figure 5a). Across all 80 samples it comprised on average 4.1% of the relative diatom sequence reads per sample, with a maximum of 76.8% of recovered amplicons on May 28^th^, 2010 (Figure S3). From the available observational data we were not able to associate this isolate with major phytoplankton blooms in Narragansett Bay. In this dataset, chlorophyll a concentrations were greater than 10 μg/L in 10 different samples. The ASV matching our *Chaetoceros* isolate was only detected above the one percent relative abundance in three of these high chlorophyll a events, but in each of these samples where it was detected it never made up more than 1.5% of the recovered amplicons.

**Figure 5:**
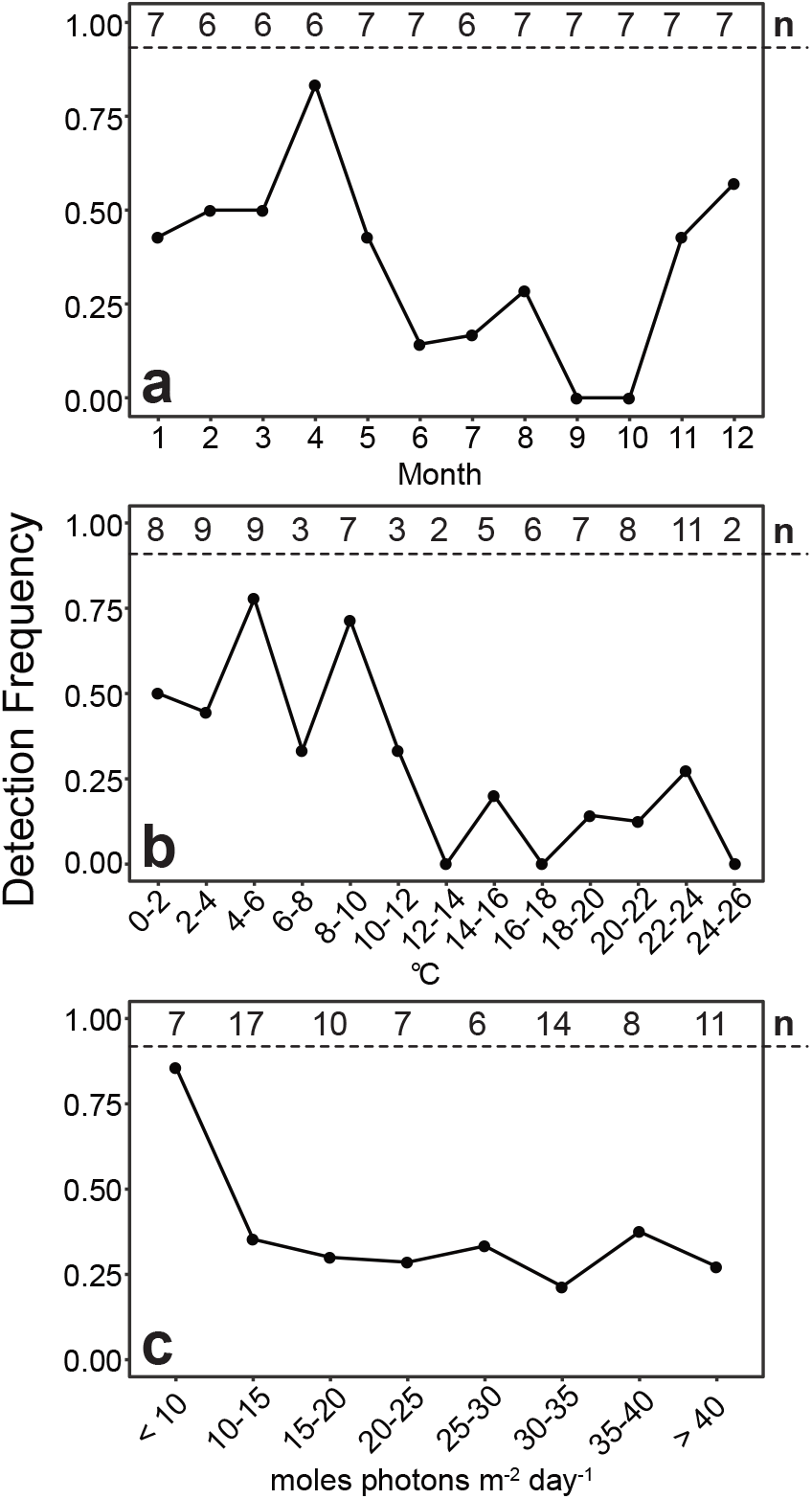
Probability of detection for the ASV matching the isolate described in this study in five years of 18S rRNA gene amplicon data by **a)** month, **b)** temperature, and **c)** seven-day average of photons received per square meter. The number of samples falling within each category is shown along the top of each graph.

To examine the thermal and light niches of the ASV matching our *Chaetoceros* isolate, we split temperatures into 13 two-degree bins from 0 to 26 °C. The isolate was present in Narraganset Bay from 0-24 deg C but occurred more frequently at temperatures < 12 °C. At these cooler temperatures the probability of observing this ASV was on average 0.52 (±0.19), but dropped to 0.11 (±0.11) above 12 °C, almost five times lower. The distribution of light exposure readings throughout the dataset was largely 10 to 40 moles photons m^-2^ day^-1^, so measurements within this range were grouped in increments of 5 moles photons m^-2^ day^-1^, with readings outside this distribution recorded as < 10 and > 40 moles photons m^-2^ day^-1^, respectively. The probability of detection was approximately equal at ~0.3 for all total irradiance levels >10 moles photons m^-2^ day^-1^; however, when total irradiance was < 10 moles photons m^-2^ day^-1^, the probability of detection nearly tripled to 0.9 (Figure 5c).

In order to assess this isolate’s geographic distribution beyond Narragansett Bay, we accessed amplicon data from the Tara Oceans dataset. Prior to analysis, a phylogenetic tree was made using the V9 region of the 18S rRNA genes used in Figure 4 to show that these primers were able to differentiate our isolate from other *Chaetoceros* spp. (Figure S4). This isolate was not detected in previously published amplicon data from the Tara Oceans project, which largely cover tropical and temperate latitudes (37). However, two ASVs that were 100% match across >90% of the sequence were detected in surface waters at 8 of 16 stations of the high-latitude Tara Oceans Polar Circle sampling (Figure 6). Both ASVs were found at the same stations with approximately equal relative abundance, and thus their results are reported together. Interestingly, they were mostly detected at stations where the relative abundance of diatom amplicons was comparatively low (Figure S5).

**Figure 6:**
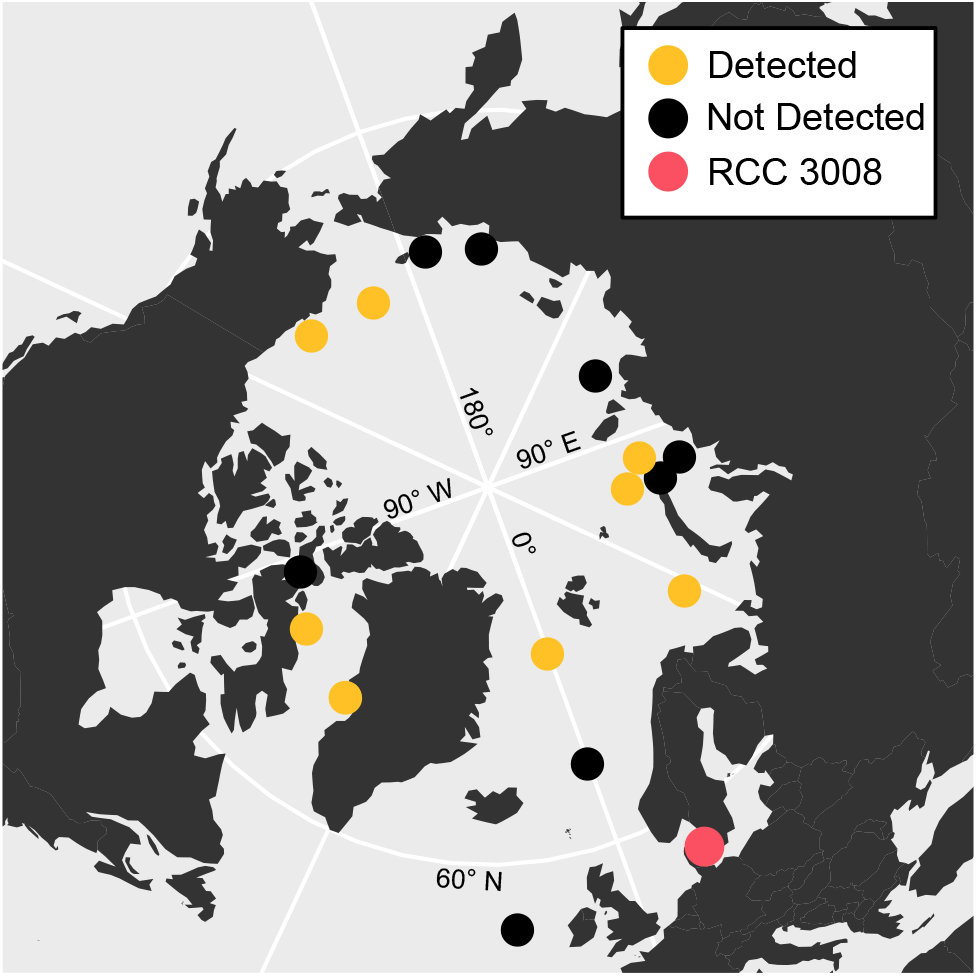
Combined relative abundance of two ASVs in the Tara Polar Ocean Circle stations that were 100% match to the V9 region of the 18S rRNA gene sequence recovered from the diatom isolate presented in this study. The red dot shows the isolation location of a diatom in the Roscoff Culture Collection whose 18S rRNA gene is a close match to this isolate.

## Discussion

In this study we describe a recently isolated nanodiatom with 18S rRNA gene sequence similarity to diatoms identified as *Chaetoceros cf. wighamii* in NCBI. In laboratory culture experiments this isolate showed a strong sensitivity to light, unusual for planktonic diatoms. This sensitivity impacted (and was impacted by) temperature, and broadly explains this diatom’s seasonal distribution across multiple years of 18S rRNA gene amplicon data from its isolation location. Despite this seemingly unusual physiology, this *Chaetoceros sp*. was detected across the samples taken during the Tara Oceans Polar Circle Expedition. This suggests its physiology may not in fact be that unusual, but rather part of a broader adaption to high latitude waters.

A preference for low light levels for growth is not necessarily uncommon in marine phytoplankton. For instance, *Prochlorococcus*, a dominant unicellular marine picocyanobacterium in the oligotrophic gyres, has well-defined low- and high-light ecotypes occupying deep and surface layers of the euphotic zone, respectively (41–43). Studies on low light *Prochlorococcus* have reported upper light limits that are similar to those we describe here for our diatom isolate (41). Even among diatoms, adaptations to low light levels have been reported in species living in benthic environments (44). However, centric diatoms are not typically considered to be part of the benthic community (although dormant resting stages can be observed)(45–47). Alternately, it could be that the low-light physiology of our *Chaetoceros* isolate is an adaptation to deeper layers of the photic zone, similar to low-light *Prochlorococcus*, but its consistent presence even in open ocean surface water samples from high latitudes does not support either conclusion. Another possibility is this diatom could be adapted to living under sea ice, where active photosynthesis can be maintained below 5 μmol photons m^-2^ sec^-1^ (48); however, this seems unlikely as temperate environments such as Narragansett Bay typically do not freeze over during the winter (49).

We also showed that at sub- or supraoptimal light levels, the maximum thermally-determined growth rate and the thermal optimum decrease. Similar observations have been made in a metanalysis of phytoplankton temperature-light effects, however at much higher light levels (50). In temperate and high latitudes, temperature and light levels increase simultaneously as spring progresses into summer and day length and angle of solar incidence increase. Our data suggest that the realized niche of this diatom is defined by the negative interactive effect of both temperature and light. For instance, in six years of amplicon data it was detected in ~45-80% of the samples taken between November and April, when temperatures are low and the days are relatively short. The frequency of detection in May could be explained by the fact that although day length is increasing, the waters in Narragansett Bay are still colder than summer and early fall conditions (mean = 15.5 ±2.7 °C). In the summer months (June through September) it was only detected 4 times. That it was detected at all during these warmer, brighter months suggests there may be additional environmental factors (such as nutrient availability) controlling its distribution. Interactions between light and temperature also explain why observations of this diatom *in situ* typically occur at temperatures well below the range of optimal growth temperatures predicted by our TPC models (13.7-17.2 °C). It should be stated however that only 5 observations were made between 10-14 °C, and it could be that more samples in this range would change the frequency of detection.

It is interesting to consider how these experimental and seasonal data may inform this diatom’s distribution in the broader ocean. For instance, although this *Chaetoceros sp*. was detected in half of the Tara Oceans Polar Circle samples, these were collected between May and October. Based on the seasonality depicted in data from Narragansett Bay this is when we would expect its abundance to be lowest. Consequently, they may be even more abundant in polar waters than observed here. SST across theses samples was low (2.56 average ±3.7 °C) compared to Narragansett Bay; however, irradiance was likely much higher due to the near constant daylight experienced during the polar summer. In culture experiments we observed that this diatom’s growth rate decreased at similar temperatures when light was supraoptimal (Figure 3a) which suggests that its physiology could be an adaptation to the cold and low-light conditions found during early spring months in the North Atlantic (51,52). Future studies using molecular methods to look at the composition of early spring bloom may show that this diatom contributes significantly to primary production at high latitudes.

Although this study documents this diatom’s singular low-light niche, more work will be needed to investigate the mechanisms involved. For photosynthetic organisms, an accumulation of deleterious reactive oxygen species (ROS) in the cell (in particular the chloroplast) is often seen following exposure to extreme irradiance (53,54). Under low light conditions, many diatoms photo-acclimate by increasing the size of their chloroplasts and the number of photosystems and antenna pigments they contain, in order to increase photon capture (55,56). It could be that our low-light *Chaetoceros* has a limited ability to adjust its photosynthetic energy acquisition systems when exposed to high light, causing a harmful buildup of ROS. Similarly, variation in the xanthophyll cycle (57,58) and production of ROS-scavenging antioxidants (58) could contribute to this planktonic diatom’s unusual physiology.

Future work should also consider the effect of light spectral quality on the irradiance and temperature interactions described here. In aquatic environments not only is the total irradiance variable, but also the availability of specific wavelengths. Shorter wavelength blue light has more energy, and thus penetrates farther into the water column than longer red wavelengths. Phytoplankton associated with low light environments such as deep water or beneath sea ice are often specialized for utilizing these higher energy wavelengths (59,60). Blue light has also been suggested to trigger germination in the resting stages of marine diatoms, and has been associated with upregulation of proteins associated with photoprotection in *Phaeodactylum tricornatum* (61,62). At higher latitudes and during the winter, solar elevation is lower compared to low latitudes or during the summer. This results in a lower angle of incidence, which causes more light to be reflected from the ocean’s surface; however, this process is skewed towards longer wavelengths, which are preferentially reflected (63). The implication is that phytoplankton at higher latitudes or during the winter season experience more blue light relative to red light. It would be interesting to test whether these diatoms experience the same light sensitivity when grown under blue light as white light (as in this study).

The interactive effects of light and temperature on this diatom’s growth in the lab and pattern of abundance *in situ* raise interesting questions about how marine phytoplankton will respond to rising temperatures associated with climate change. It is broadly suggested that organisms at high latitudes exist at temperatures well below their thermal optima, and therefore rising temperatures will be advantageous, increasing their growth rate (23,64). The average temperature at Narragansett Bay in the five years of temperature data accompanying this amplicon dataset is 12.4 °C, below the optimal temperatures predicted by our three TPC models. However, because of the strong regulation of thermal niche by light level in this *Chaetoceros sp*. it could be that this isolate will not fare better with rising temperatures, as warmer conditions increase its susceptibility to light stress. In a shallow (8m), well mixed estuary such as Narragansett Bay this interaction between light and temperature may in fact shrink the range of months where growth of this diatom is feasible. For instance, it is frequently (appearing in > 40% of the samples) observed as late in the year as April and May, where day length is longer and solar elevation higher than during the winter months. Rising temperatures during those months may be harmful, increasing the diatom’s susceptibility to light stress; although it may also be that rising temperatures will be advantageous during winter months (e.g. December to March) when light levels are seasonally low.

In the open ocean, this diatom’s sensitivity to light may disadvantage it in a warmer future. Current models predict a shoaling of the thermocline at high latitudes, increasing light exposure by trapping phytoplankton closer to the surface (65). This is expected to increase overall photosynthetic growth, as high latitude phytoplankton are often considered light limited (66); however, the *Chaetoceros* isolate described here which was observed across the polar circle challenges this paradigm. Future work at high latitudes should further investigate the abundance of this diatom (especially during early spring bloom conditions) in order to better predict how climate change will impact phytoplankton communities in these regions.

Our study highlights one facet of the largely unrecognized but almost limitless diversity that exists in marine phytoplankton communities. It is fascinating that a planktonic diatom with such a specialized light and temperature niche was discovered at the longest running phytoplankton timeseries in existence, and that despite its specialization it is one of the more common diatoms at this well-studied site. This work also shows that light and temperature can interact to define a thermal niche. Even in species that thrive at comparatively high light levels, changes in light could similarly impact their response to changes in temperature and influence how they will fare in a warming ocean. Future studies should consider high light as an interactive variable along with other co-stressors such as elevated temperatures and CO_2_ when predicting how phytoplankton will respond to global change.

## Acknowledgements

We would like to thank Amanda Montalbano, Meghan Phan and Roxanna Andrade for assistance with preliminary culture work and Daniel Campo for help with library prep and Illumina sequencing. Funding was provided by National Science Foundation (NSF) grants OCE1538525 and OCE1638804 to DAH, and OCE1638834 to TAR. Part of this research was conducted using the University of Rhode Island’s Marine Science Research Facility, supported by NSF EPSCoR awards 1004057 and 1655221.

