## Supplemental Figures for "Irradiance modulates thermal niche in a previously undescribed low-light and cold-adapted nano-diatom"

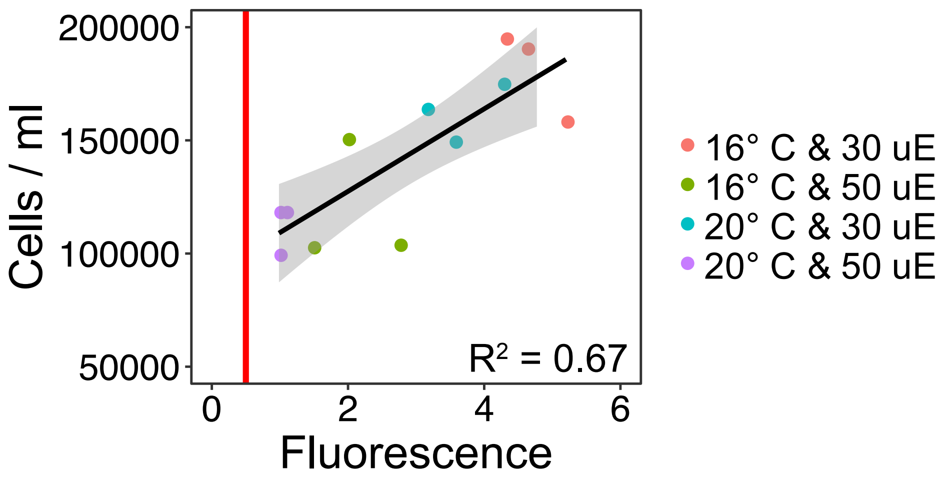


Figure S1: Comparison of fluorescence across cell concentrations for cultures in exponential phase for held at either 16 or 20 °C or 30 or 50 µmoles photons $m^{-2}$ $sec^{-1}$. Vertical red line shows the fluorescence all cultures were diluted to at the beginning of each semi-continuous growth cycle. Black line is the result of a simple linear regression with the $r^2$ shown.

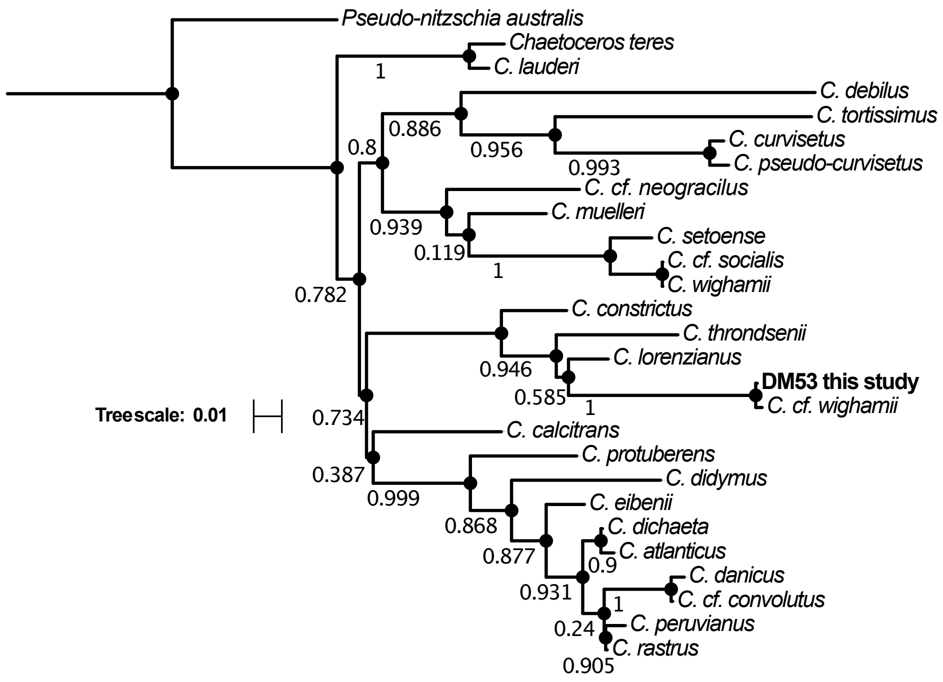


Figure S2: Maximum likelihood tree for the genus *Chaetoceros* using the V4 hypervariable region of the 18S rRNA gene.

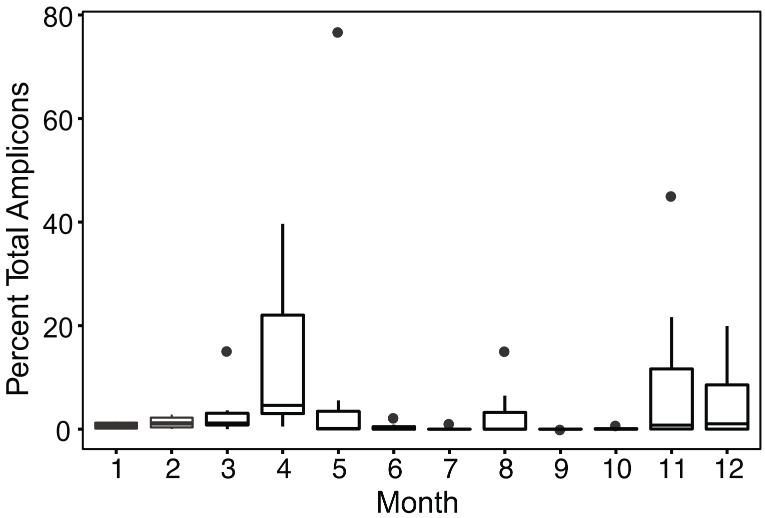


Figure S3: Relative abundance of ASV6 (recovered ASV matching the 18S rRNA sequence from our isolate) for each month throughout six years of monthly sampling.

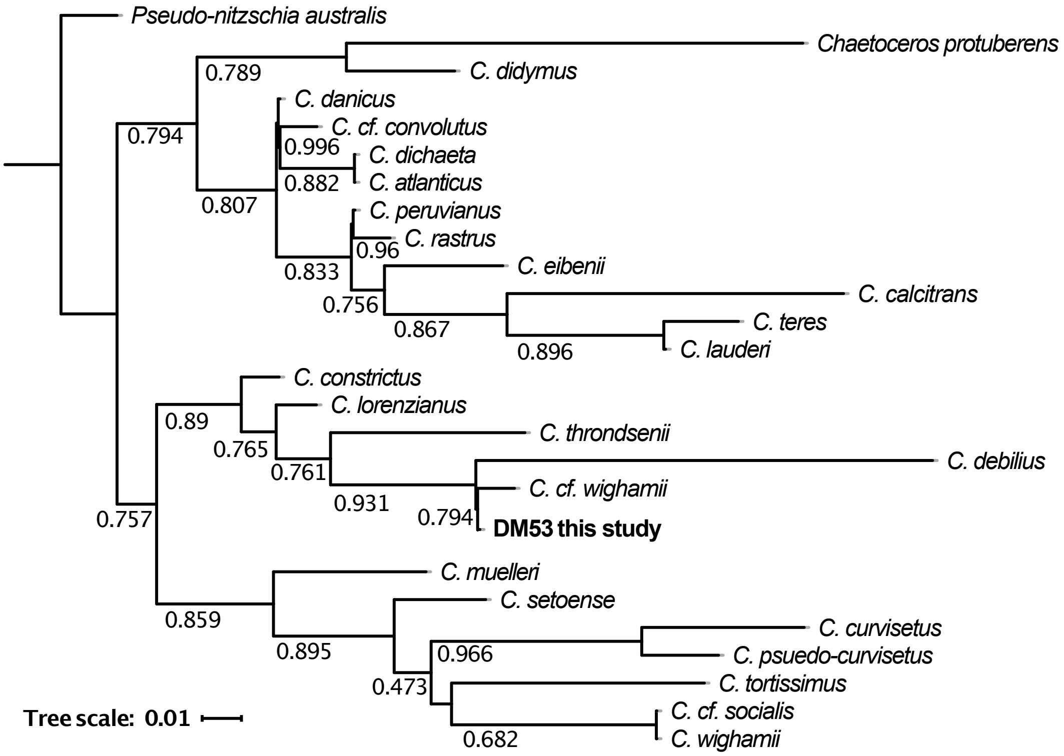


Figure S4: Maximum likelihood tree for the genus *Chaetoceros* using the V9 hypervariable region of the 18S rRNA gene.

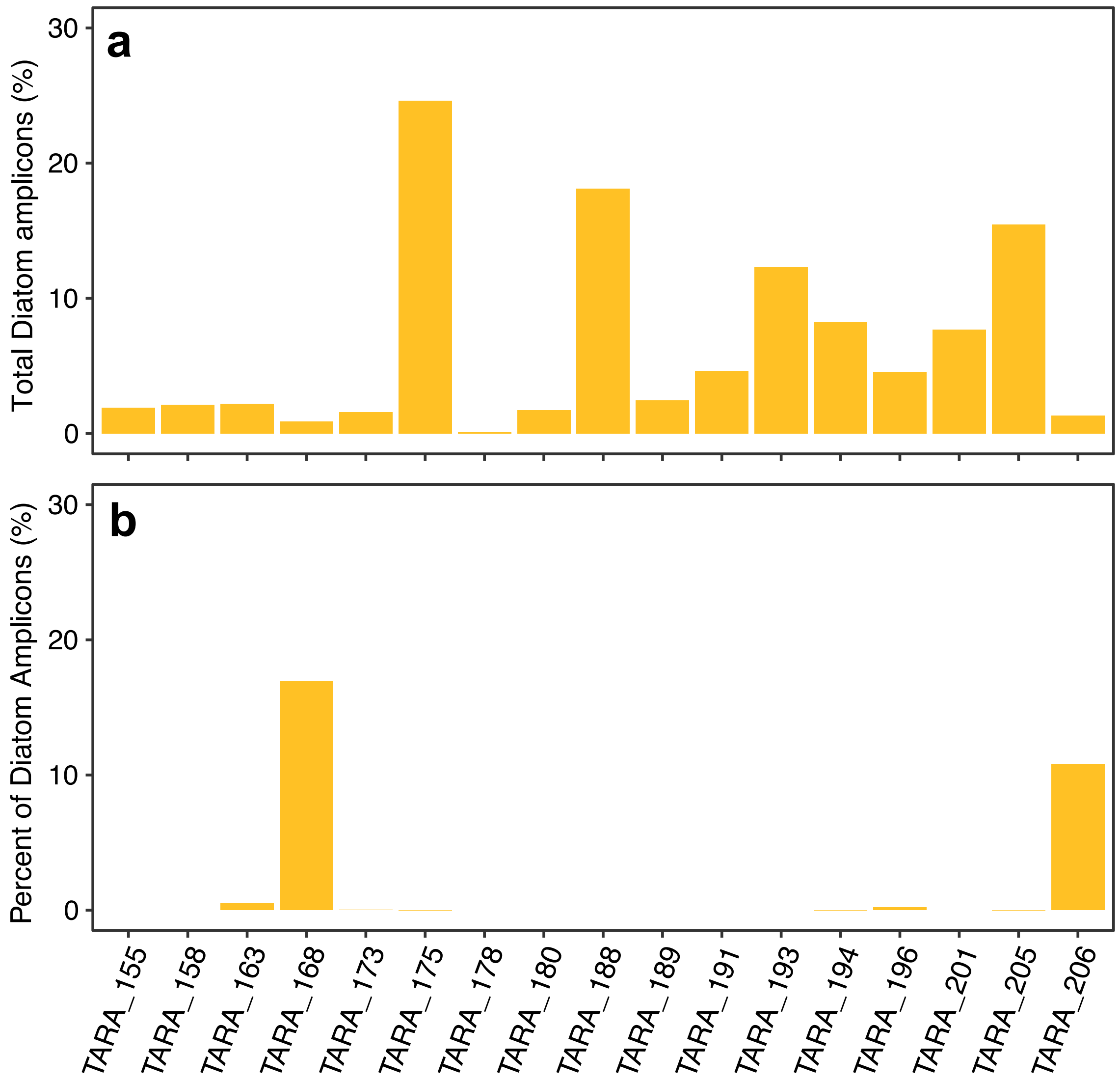


Figure S5: **a)** The percentage of recovered V9 amplicons that belonged to diatoms, and **b)** the percentage of all diatom amplicons that matched the *Chaetoceros sp.* isolate described in this study.
